# Proteome of actin stress fibers

**DOI:** 10.1101/2021.06.01.446528

**Authors:** Shiyou Liu, Tsubasa S. Matsui, Na Kang, Shinji Deguchi

## Abstract

Stress fibers (SFs), which are actomyosin structures, reorganize in response to various cues to maintain cellular homeostasis. Currently, the protein components of SFs are only partially identified, limiting our understanding of their responses. Here we isolate SFs from huma fibroblasts HFF-1 to determine with proteomic analysis the whole protein components and how they change with replicative senescence (RS), a state where cells decline in ability to replicate after repeated divisions. We found that at least 263 proteins are associated with SFs, and 101 of them are upregulated with RS, by which SFs become larger in size. Among them, we focused on eEF2 (eukaryotic translation elongation factor 2) as it exhibited upon RS the most significant increase in abundance. We show that eEF2 is critical to the reorganization and stabilization of SFs in senescent fibroblasts. Our findings provide a novel molecular basis for SFs to be reinforced to resist cellular senescence.

## Introduction

Stress fibers (SFs) are bundles of actin filaments expressed in mesenchymal cell types such as fibroblasts (Kreis and Birchmeier, 1980). SFs exhibit adaptive responses to changes in intracellular and extracellular cues by modulating the assembly and disassembly (Tojkander *et al*., 2012), which allows them to be involved in diverse cellular functions. Typical examples include the regulation of cell–substrate adhesion (Braga *et al*., 1997; Livne and Geiger, 2016), cell migration (Hotulainen and Lappalainen, 2006), mechanotransduction (Chen *et al*., 2004; Burridge and Wittchen, 2013; Hirata *et al*., 2015; Wei *et al*., 2020), morphological maintenance (Meng and Takeichi, 2009; Jalal *et al*., 2019; Kang *et al*., 2020), mechanical properties (Deguchi *et al*., 2006; Kumar *et al*., 2006), and avoidance of proinflammatory signaling and resulting cell homeostasis (Kaunas *et al*., 2006; Chien, 2007; Kaunas and Deguchi, 2011).

Currently, the components that constitute SFs remain incompletely identified. In fact, only 20 proteins were listed in a recent review to be associated with the complex of SFs, which include cytoskeleton-binding proteins, adhesion proteins, and small GTPase-related enzymes (Tojkander *et al*., 2012). Given, however, that more than 900 proteins have been reported based on proteomic analysis as constituting focal adhesions (FAs) connected to the termini of individual SFs to work cooperatively (Kuo *et al*., 2011) and that SFs are engaged in such diverse biological functions under multifarious stimuli, it is likely that SFs actually consist of a much larger number of proteins. Thus, here we aim at comprehensively identifying the protein components inherent to SFs and how they change as cells undergo senescence, which is also a biological process that can intricately be linked to age-related diseases.

Replicative senescence (RS) is defined as a state in which normal somatic cells including fibroblasts decline in their ability to replicate in vitro after a long period of culture and repeated divisions (Goldstein, 1990; Muller, 2009). This cellular aging leads to irreversible growth arrest accompanied by phenotypic changes such as flat cell morphology and increased senescence-associated β-galactosidase (SA-β-gal) activity. The close association with the cell morphology suggests the involvement of SFs in RS. In fact, there are many reports describing their morphological change associated with RS, although the response seems to vary depending on the cell types (Hwang *et al*., 2009). Meanwhile, RS-driven changes in the molecular components of SFs remains poorly understood except for actin, i.e., only their major constituent.

Here we investigate the effect of RS on SFs in human fibroblasts, typically used in RS-related studies. To this end, we isolate SFs from two different cell populations, young or aged, to compare the morphology and protein components between them with proteomic analysis. We found that SFs, identified here to consist of at least 263 different types of proteins, become thickened upon RS with increased expression of 101 proteins including major proteins NMMIIa and α-actinin-1. Among them, we focused on a specific protein eEF2 (eukaryotic translation elongation factor 2) as it exhibited upon the induced senescence the most significant increase in abundance in SFs while categorized as having diverse biological functions in the proteomic analysis. We then show that eEF2 is critical to the reorganization of SFs that are stabilized in senescent cells where proliferation and migration are both reduced. eEF2 is well characterized as an essential factor for promoting the mRNA translation by ribosomes (Choi *et al*., 2018), but has rarely been reported with respect to the cytoskeleton. Thus, our findings provide a novel molecular basis for SFs to be reinforced to resist fibroblast senescence.

## Results

### RS induces thicker SFs in fibroblasts

We induced RS to human foreskin fibroblasts HFF-1 by repeated passaging. The degree of the senescence was identified using several indicators, where the increases in SA-β-gal activity, p21 expression, and p53 expression are among the most common indicators. We found significant upregulation of these indicators upon the increased passage numbers, in which passage number 2, 10, and 33 (termed P2, P10, and P33, respectively) were selected as the target of the present analysis (Fig. 1A,1B). P2, which has the relatively lowest degree of senescence, is regarded as a young control for the subsequent analyses. We compared the phenotypic characteristics, in which more senescent cells exhibited slower migration speed (Fig. 1C), lower proliferation (Fig. 1D), and increased cellular and nuclear areas (Fig. S1). Focusing on individual SFs observed with phalloidin staining, they exhibited thicker morphology as RS proceeded, while there was no significant difference in the number per cell (Fig. 2).

**Figure 1:**
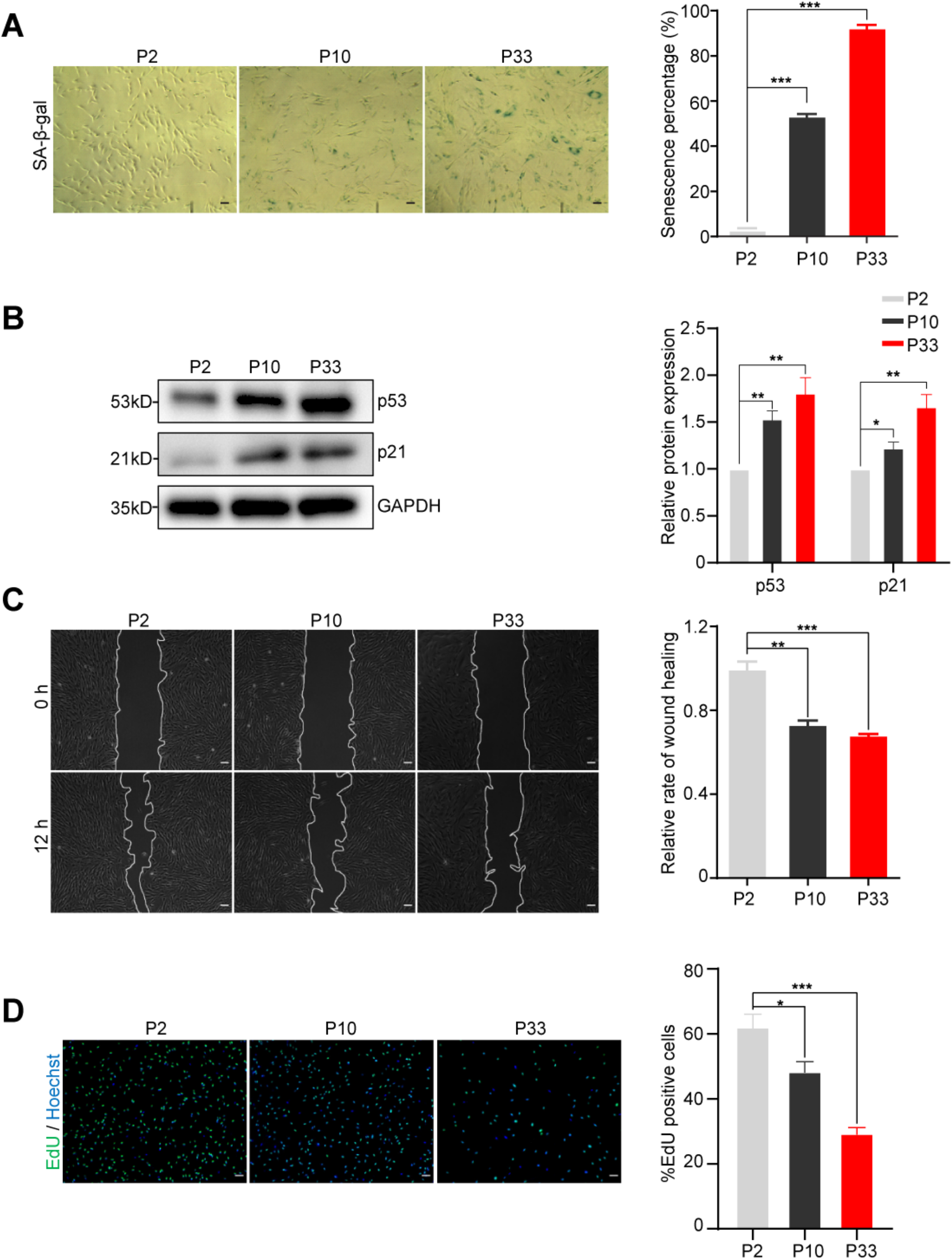
Induction of RS on HFF-1 cells. (A) The extent of senescence in cells passaged 2, 10, and 33 times (shown as P2, P10, and P30, respectively) was evaluated 24 hours after SA-β-gal staining and quantified. (B) Protein expression levels of p53 and p21 were analyzed with western blotting and quantified. (C) Wound healing assay was performed for 12 hours and quantified. White curves in the images represent the front border of the migrating cells. (D) EdU proliferation assay was performed for 36 hours and quantified. Scale, 100 μm.

**Figure 2:**
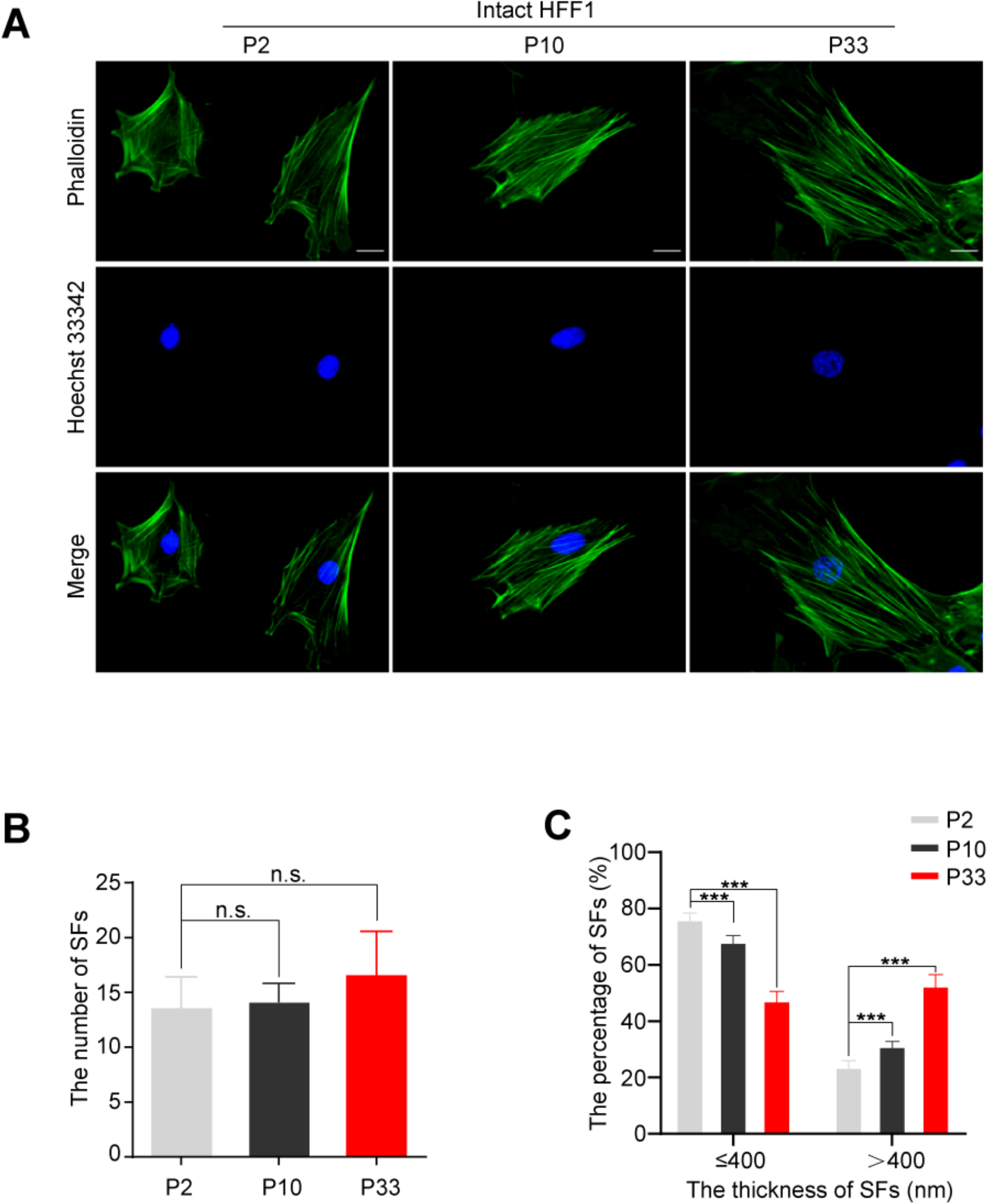
RS induces thick SFs. (A) Actin filaments (phalloidin) and nuclei (Hoechst) in HFF-1 cells at different senescence levels of P2, P10, and P33. (B) The number of individual SFs per cell (*n* = 20 cells from 3 independent experiments). (C) The ratio of the number of individual SFs in cells with a thickness of ≦ 400 μm to that of > 400 μm (*n* = 20 cells from 3 independent experiments). Scale, 100 μm.

### Validation of the isolation of SFs from cells

Motivated by the above finding that the morphology of SFs is altered upon RS, we attempted to comprehensively identify the protein components of SFs that markedly change with RS. To this end, SFs were isolated from the rest of the cells for proteomic analysis (Fig. 3A). Specifically, cells were treated with hypotonic shock to finally remove most of the cell bodies according to our previously developed techniques (Matsui *et al*., 2011; Okamoto *et al*., 2020), leaving ventral SFs physically attaching to the substrate. These SFs remained unchanged in appearance during the hypotonic shock treatment as observed with YFP-actin transiently expressed in the cells (Fig. S2A). The ventral SFs, now exposed to the solution, were next treated with a gentle detergent (0.05% Triton X-100) in a cytoskeleton stabilizing buffer to remove the underlying plasma membrane and its associated proteins as well as the nucleus. Factin staining (Fig. 3B) and immunofluorescence of anti-α-actinin-1, nonmuscle myosin IIa (NMMIIa), and β-actin (Fig. 3C) showed that these proteins remain associated along the length of the extracted SFs even after the cell bodies are removed. The RS-elicited increase in the thickness of individual SFs, observed in intact cells (Fig. 2), was also detected even after they were extracted from the cells (Fig. S2B, S2C). The presence of these proteins, in addition to nonmuscle myosin regulatory light chain (MLC), was also confirmed in the extracted SFs with western blotting performed by physically isolating them from the dish using a rubber scraper (Fig. 3D). Meanwhile, other proteins that are not typically associated with SFs – specifically, vinculin, tubulin, and GAPDH – were almost absent in the isolated SF lysates, but instead they were detected in the cell body lysates. These results support that subcellular materials that inherently constitute SFs were indeed isolated from the rest of the cells while keeping the apparent morphology and compositions.

**Figure 3:**
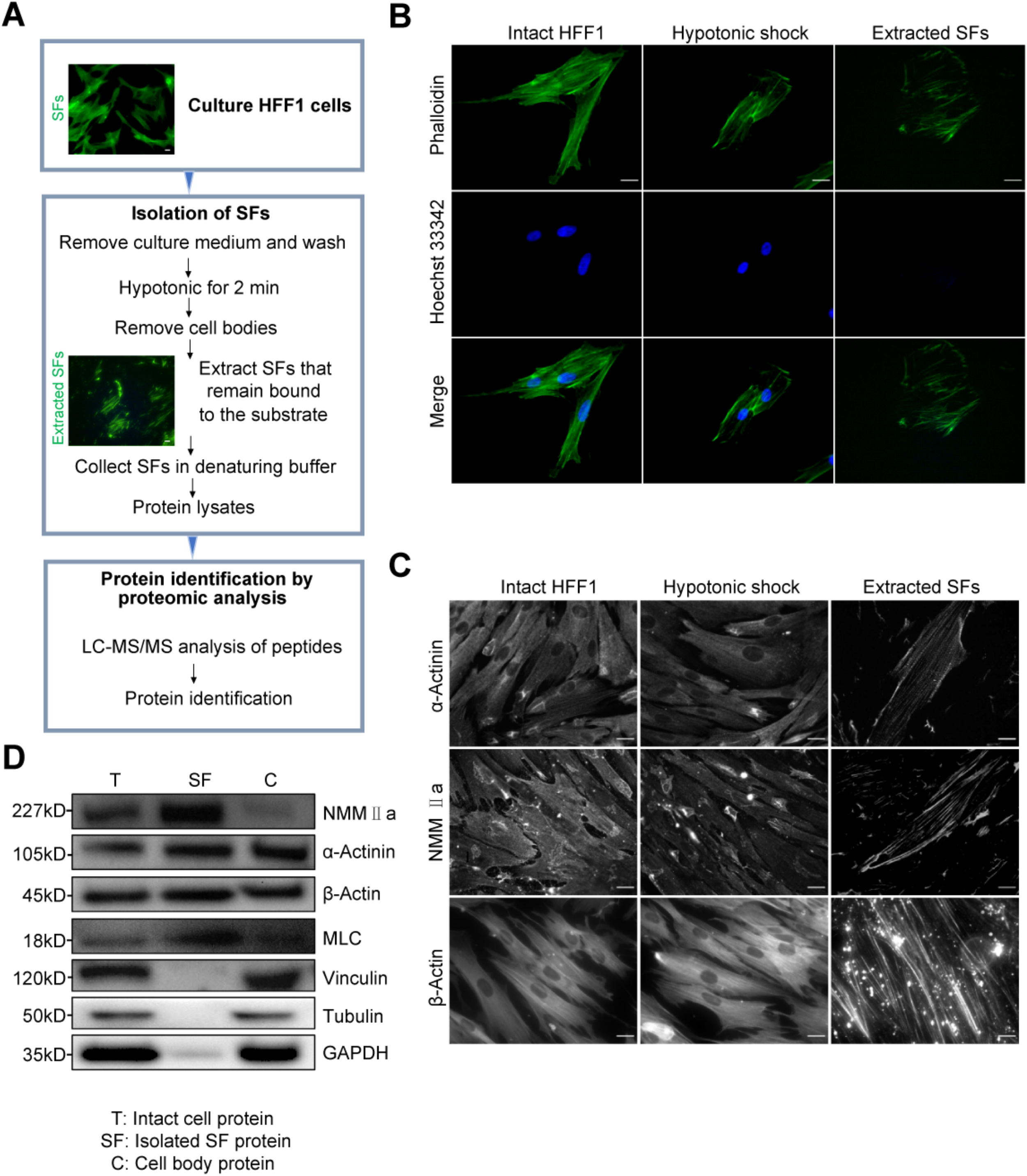
Isolation of SFs. (A) Flow chart for the isolation of SFs from HFF-1 cells and the component identification by proteomic analysis. Intact HFF-1 cells (top panel) and isolated SFs (middle panel) were stained with phalloidin. (B, C) Actin filaments (phalloidin) and nuclei (Hoechst) (B) and immunofluorescence of α-actinin-1, NMMII a, and β-actin (C) in intact HFF-1 cells at P2 (left), those subjected to hypotonic shock (middle), and those after the extraction procedure (right). Note that the nuclei are finally removed and thus are absent in the extracted SF samples. (D) Immunoblot of the proteins contained in total cell lysates (T), isolated SF lysates (SF), and cell body lysates (C). Scale, 20 μm.

### Proteomic analysis reveals RS-induced upregulation of eEF2 within SFs

We performed shotgun proteomic analysis on the isolated SFs to evaluate how they change in composition between the two extremely different passage numbers P2 and P33 (Fig. 3A). We identified that at least 263 different proteins are associated with the isolated SFs (data not shown): the detected proteins are categorized to 3 groups in accordance with the magnitude of the fold change (FC) in the ratio of expression level of each protein at P33 to that at P2 as FC > 2, 2 ≥ FC ≥ 0.5, or 0.5 > FC, i.e., representing that, in response to RS, the protein is upregulated, almost unchanged, or downregulated, respectively. The number of the detected proteins, 263, is much larger than that previously listed as the components of SFs, 20 (Tojkander *et al*., 2012), thus potentially allowing us to detect novel proteins that were not known to be associated with SFs. Of the 20 proteins, on the other hand, only 13 were identified in our study (specifically, MYH9, PALLD, SEPT2, FLNA, TPM4, CALD1, PDLIM7, CSRP1, ACTN1, VASP, CNN2, ZYX, and TAGLN), but the rest was not detected for some reason, e.g., the specific cell types used, protein sensitivity to different experimental treatments, and/or small amount of the protein association to SFs.

Regarding all the 263 proteins, we performed a literature/database review (particularly using GeneCards (Weizmann Institute of Science)) to identify ones that have been well documented to be related to the cytoskeleton. Consequently, 58 are listed to be associated with actin filaments and/or SFs, and 15 are with intermediate filaments and/or microtubules (Fig. S3). Aiming at identifying novel SF-associated proteins, here we focused on the remaining ones, i.e., 190 (= 263 - 58 −15), for the subsequent analysis. These proteins are categorized in accordance with the known biological processes using Scaffold software (Proteome Software) into one or more of the following 8 groups (Fig. 4): “Biological Regulation” (74), “Cellular Process” (135), “Developmental Process” (43), “Establishment of Localization” (42), “Immune System Process” (20), “Metabolic Process” (98), “Multicellular Organismal Process” (42), and “Response to Stimulus” (37), in which the figure in parenthesis represents the number of proteins included in the respective group. As RS is related to diverse biological activities such as development, metabolism, stress, and cancer progression (van Deursen, 2014), it may not be surprising that the 190 SF-associated proteins range over such various categories.

**Figure 4:**
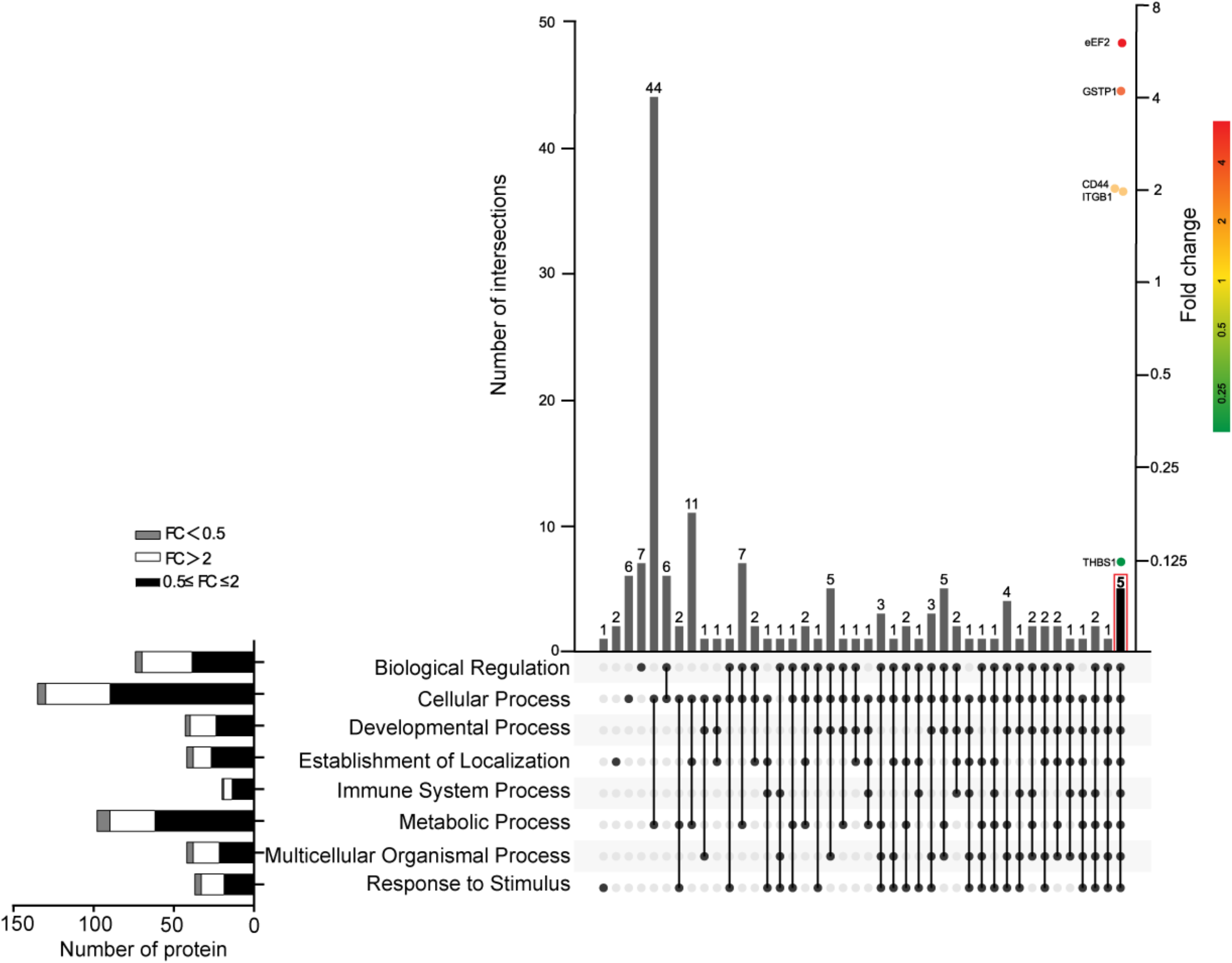
Proteomic analysis of isolated SFs reveals significant increase in eEF2 expression upon RS. In total 263 proteins were identified to be associated with the isolated SFs. Among them, 73 were already commonly known to be associated with the cytoskeleton according to the GeneCards database (Fig. S3), and therefore the rest (190 proteins) were focused on here and categorized according to the known 8 biological processes (described below in the diagram) using the Scaffold software. Of these, eEF2 is one of the 5 proteins involved in all the 8 processes and shows the most significant increase, among the 5, in the amount of the association to SFs in response to RS.

Next, we sought for proteins that exhibited distinct responses with an FC of >2 or <0.5, as well as that are implicated in all the above 8 biological processes. Consequently, we obtained the following 5 proteins: THBS1, CD44, ITGB1, GSTP1, and eEF2 (Fig. 4). Among them, eEF2 was the most significantly upregulated protein with an FC of 6.1. Taken together, we isolated SFs from fibroblasts with or without RS and found that approximately 70% (190 out of 263) of the associated proteins had not been well characterized as the constituents of SFs. In view of the high sensitivity to RS, relevance to diverse biological processes, and lack in understanding of its involvement in SFs, hereafter we focus on the specific protein eEF2.

### Colocalization of endogenous eEF2 with SFs becomes prominent upon RS

Consistent with the above proteomic analysis, western blotting on the isolated SFs showed that the expression in eEF2, known as one of the elongation factors, increases with the progress in RS (Fig. 5A). Anti-eEF2 immunofluorescence was performed to investigate the localization of endogenous eEF2 in HFF-1. For young cells at P2, eEF2 is mainly localized in the cytoplasm but is also weakly detected along the length of SFs (Fig. 5B, 5C). At P33, the colocalization becomes prominent as eEF2 shows fibrous patterns like those of SFs. The extent of the accumulation to SFs was quantified by taking the ratio of eEF2 intensity at P10 or P33 to that at P2, showing that it significantly rises as the senescence proceeds to be more than twice at P33 compared to P2 (Fig. 5D). We observed a similar RS-induced eEF2 upregulation with western blotting, which occurs along with an increased expression in NMMIIa, α-actinin-1, and β-actin, i.e., representative proteins associated with SFs (Fig. 5E, 5F).

**Figure 5:**
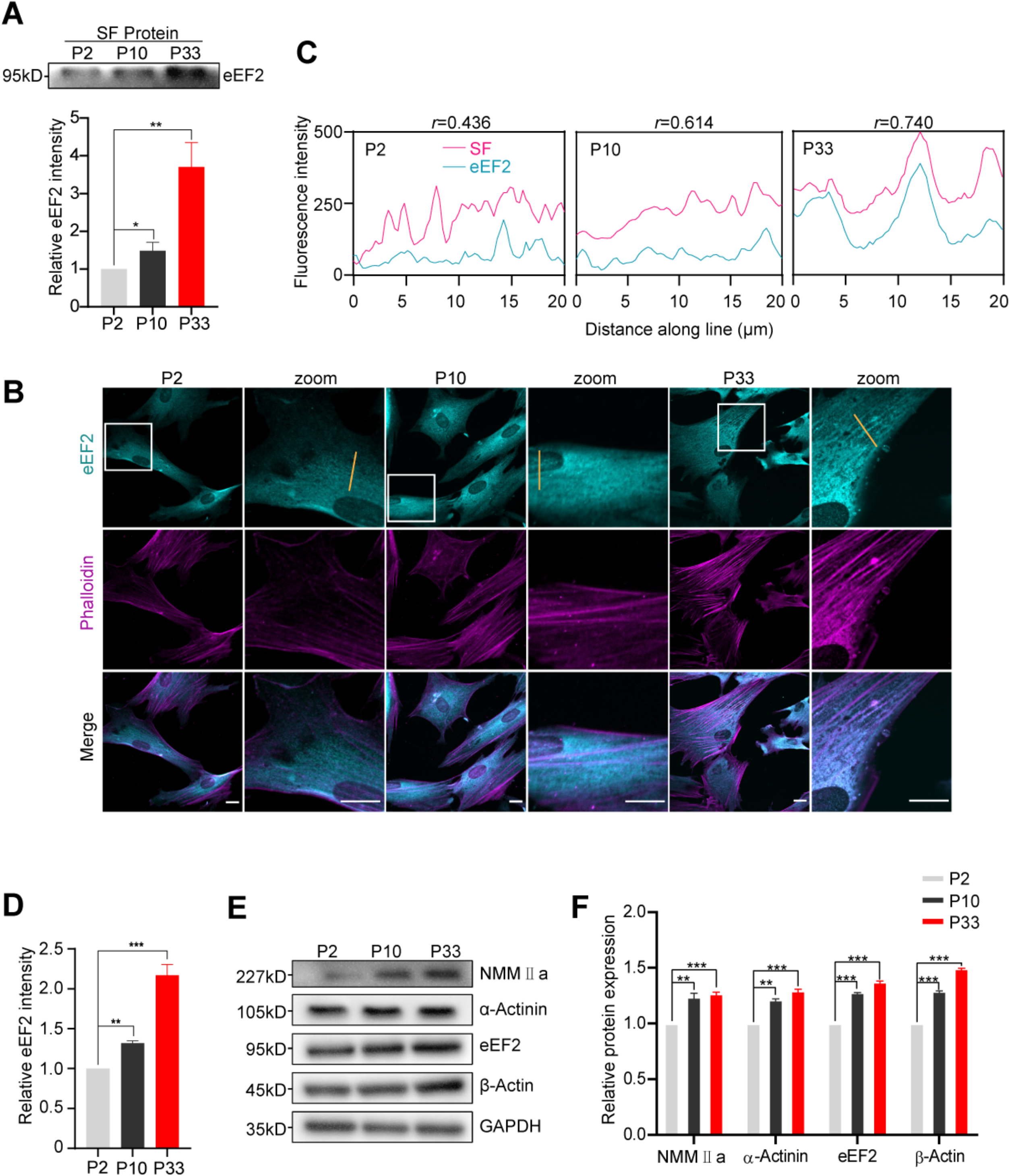
RS enhances localization of eEF2 to SFs. (A) Western blot shows increased expression of eEF2 in isolated SFs upon RS. (B) Immunofluorescence of eEF2 to observe the endogenous localization at the different senescence levels. The areas with white rectangles are magnified on the right in each case. (C) Intensity distributions along the length of the orange lines in B show increased localization of eEF2 on SFs (phalloidin) with a Pearson’s coefficient *r* of up to 0.740 at P33. (D) Quantification of the ratio of eEF2 fluorescence intensity at P10 or P33 to that at P2. (E, F) Western blot shows increased expression in eEF2 and major SF proteins in intact cells in response to RS. Scale, 20 μm.

### eEF2 is critical to the RS-induced enhancement of SFs

To explore if eEF2 has a role in the RS-mediated enlargement of SFs, or the increased expression of eEF2 is just a consequence of the enlargement of SFs, we silenced eEF2 using siRNA in HFF-1 at P33 and observed the response of SFs. The immunofluorescence showed that the abundance of endogenous eEF2 is decreased by the silencing, and consequently SFs particularly located in the cytoplasm are significantly weakened or disappear (Fig. 6A). The remaining SFs under eEF2 silencing were significantly decreased in thickness (Fig. 6B). The expression of NMMIIa and α-actinin-1 was significantly downregulated upon eEF2 silencing, but that of β-actin was not changed significantly (Fig. 6C, 6D).

**Figure 6:**
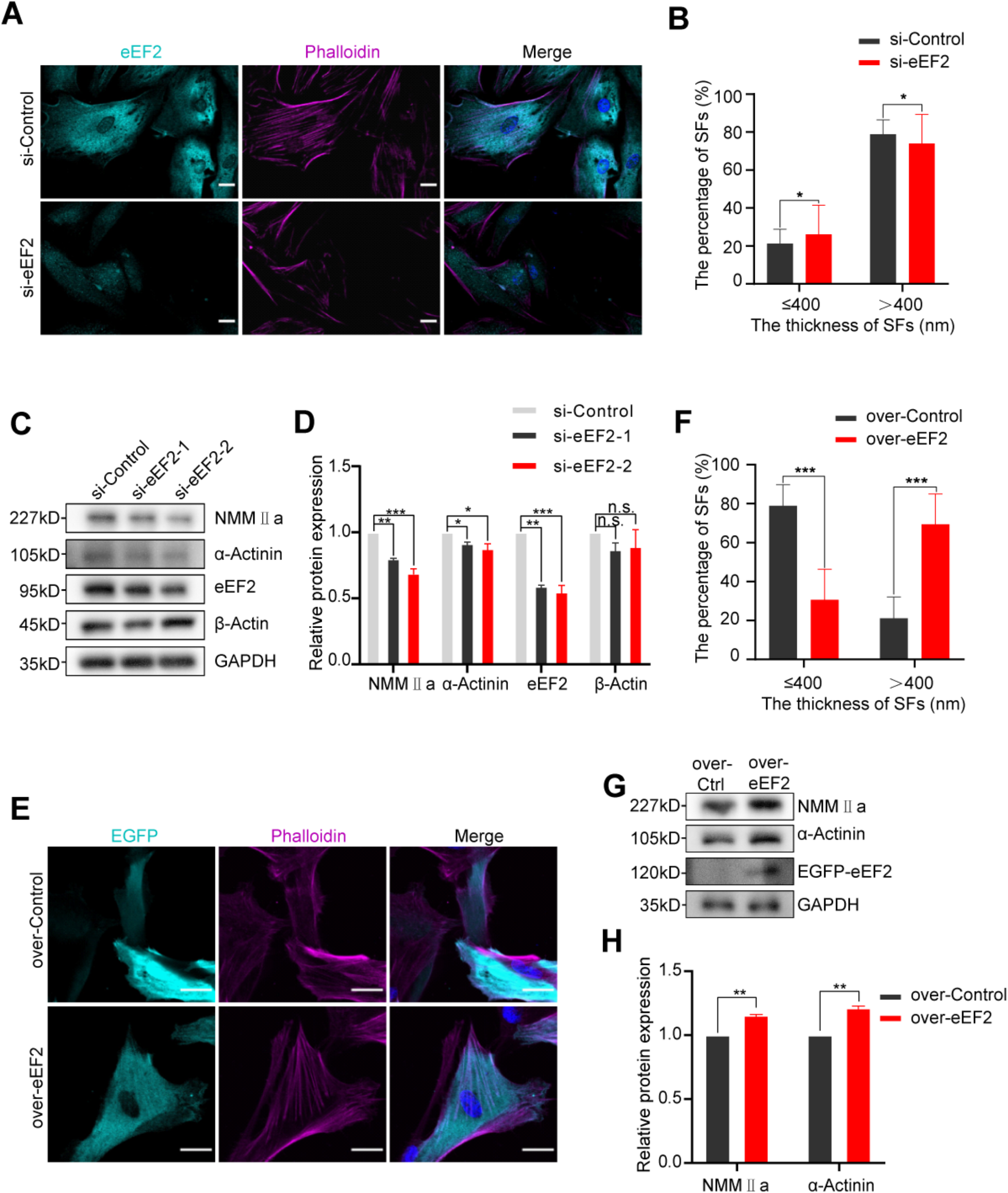
eEF2 is critical to SFs. (A) Immunofluorescence of eEF2 and F-actin staining in cells at P33 expressing siRNA for control or for eEF2. (B) Quantification shows that the thickness of individual SFs in the cytoplasm is decreased upon the silencing of eEF2. (C, D) Western blot shows decreased expression in NMMIIa and α-actinin-1 upon the silencing of eEF2 (where two types of siRNAs were used), but the concomitant change in β-actin is not significant. (E) Immunofluorescence of EGFP and F-actin staining in cells expressing EGFP for control or EGFP-eEF2. (F) Quantification shows that the thickness of individual SFs is increased upon the overexpression of eEF2. (G, H) Western blot shows increased expression in NMMIIa and α-actinin-1 upon the overexpression of eEF2. EGFP-eEF2 was detected by the eEF2 antibody. Scale, 20 μm.

We also analyzed the effect of overexpressing eEF2 in young HFF-1 at P2 to find that more visible, thicker SFs appear upon the overexpression compared to control (Fig. 6E, 6F). The consistent tendency was observed with western blotting, in which the expression of NMMIIa and α-actinin-1 is significantly upregulated upon eEF2 overexpression (Fig. 6G, 6H).

We next analyzed the effect of eEF2 on the stability of SFs by performing fluorescence recovery after photobleaching (FRAP) experiments. EGFP-tagged MLC was used as a marker of SFs in the senescent HFF-1 at P33 with or without eEF2 silencing. The fluorescence of EGFP-MLC recovered faster with the silencing, suggesting that the depletion of eEF2 destabilizes the senescent SFs (Fig. 7A, 7B). This tendency was quantified by analyzing the mobile fraction of the fluorescence, which measures the instability of SFs and was indeed found to significantly increase with eEF2 silencing (Fig. 7C). Together, these results suggest that eEF2 mediates and stabilizes the RS-induced enhancement of SFs.

**Figure 7:**
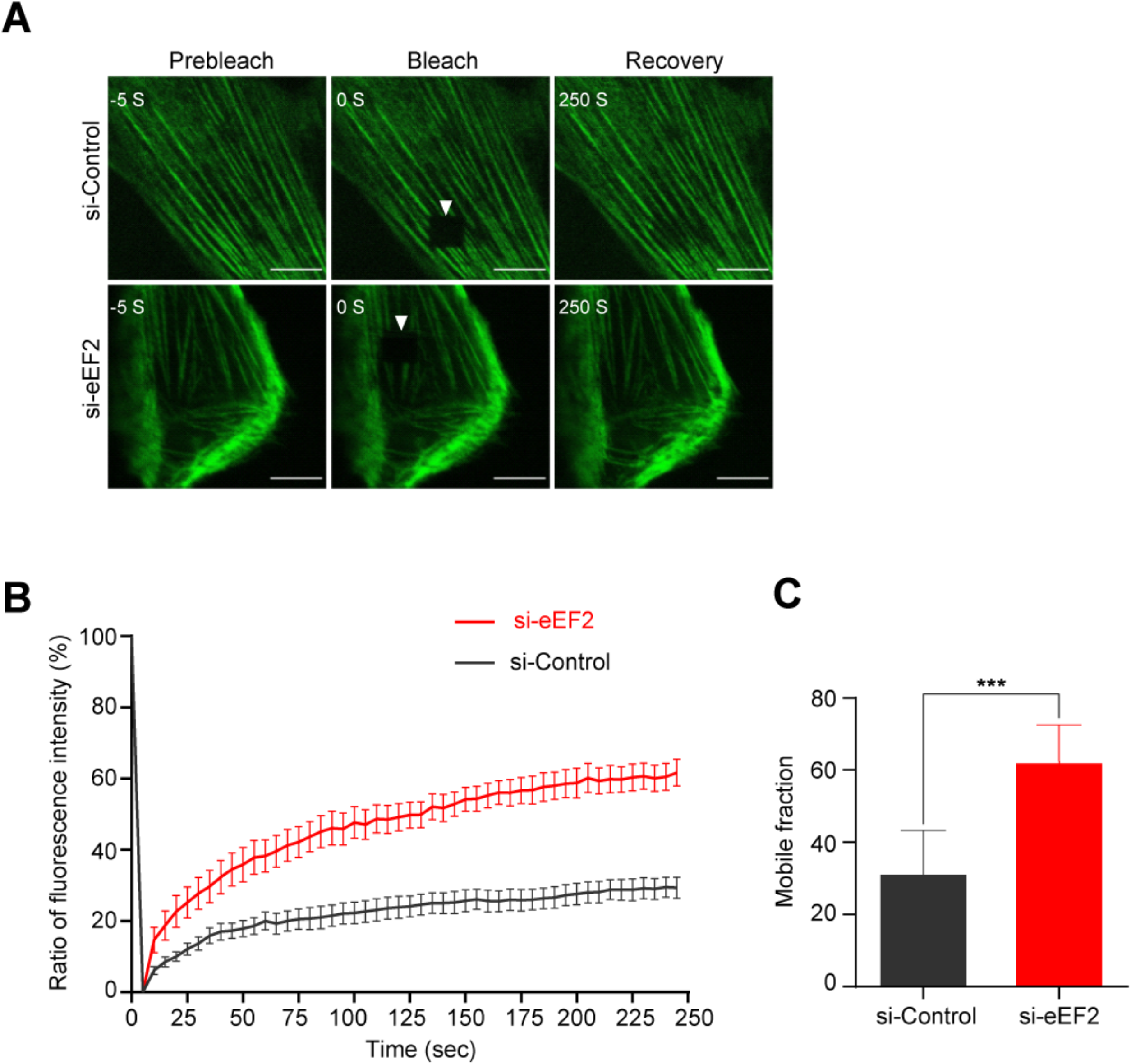
eEF2 stabilizes SFs. (A) FRAP experiments were performed on control cells expressing EGFP-MLC and those cells with eEF2 silencing. A rectangular region (indicated by the arrowhead) was bleached at time zero. (B) Time-series change in fluorescence intensity (mean ± standard deviation from 3 independent experiments), in which all the values are normalized between the initial and photobleached states and shown in percentage. (C) Quantification of mobile fraction shows that eEF2 silencing results in destabilizing the EGFP-MLC-labeled SFs. Scale, 10 μm.

## Discussion

Human fibroblasts have been used for decades as a model for understanding the basis of cellular senescence (Goldstein, 1990). Particularly, RS of fibroblasts is known to capture the features of cellular senescence in tissues (Hayflick, 1965; Cristofalo *et al*., 2004). Indeed, we observed increased SA-β-gal positive cells and elevated expression of p53 and p21 in human fibroblasts HFF-1, which are recognized as reliable markers of cellular senescence (Levine and Oren, 2009; McHugh and Gil, 2018). Using this cell line, we demonstrated that proliferation, migration, and enlargement of cellular and nuclear areas are all altered upon RS in a manner typical of the actual senescence.

Along with these responses, interestingly, a significant thickening of individual SFs is induced in senescent cells. It remains, however, controversial whether senescence leads to thicker or thinner SFs (Chen *et al*., 2000; Nishio and Inoue, 2005), despite their inherent connection to cell morphology and migration (Hotulainen and Lappalainen, 2006; Meng and Takeichi, 2009; Lin *et al*., 2017; Kang *et al*., 2020; Kang *et al*., 2021). As fibroblasts in tissues are subjected to various environmental cues including hormone signaling and wound remodeling, these individual differences are likely to be caused by complicated mechanisms (Hwang *et al*., 2009). Nevertheless, to gain better insight into the mechanisms, not only just observing the morphology, but also specifying the molecular identity intrinsic to SFs at different senescence stages must be indispensable. We therefore attempted to physically isolate individual SFs from HFF-1 and explore their protein reorganization in response to RS. We then found that there is indeed a significant compositional change upon RS.

To our knowledge, the present study is the first that comprehensively elucidates the proteome of SFs and furthermore how it changes upon cellular senescence. Despite increasing research on SFs, the knowledge on their components had been highly limited (Tojkander *et al*., 2012). This situation on SFs was distinct from that on FAs as a comprehensive proteomic analysis was previously performed on isolated FAs to eventually identify that 905 proteins are associated with FAs, while 459 of them change in abundance upon myosin II inhibition (Kuo *et al*., 2011). The isolation of FAs was achieved with hypotonic shock and subsequent strong trituration on HFF-1 fibroblasts while minimizing contamination by SFs. In the present study, we took a similar approach by isolating SFs with hypotonic shock and more gentle fluid shearing forces on the same fibroblast cell line. This isolation method using a cytoskeleton-stabilizing buffer has been validated in many aspects including the maintained helical microstructures as well as contractile function (Matsui *et al*., 2011; Deguchi *et al*., 2012; Okamoto *et al*., 2020). We consequently succeeded in specifying at least 263 proteins as the components of SFs, and among them 101 and 7 are upregulated and downregulated upon RS, respectively. These large numbers seem to reflect the variety of biological processes to which SFs are potentially related, and thus given the lack of relevant proteome information the data obtained here will be of value in further studies.

Of the 263 proteins, RS increases approximately 38% (≈ 101/263*100) in expression abundance, while only less than 3% (≈ 7/263*100) is decreased, suggesting that SFs are reinforced to resist cellular senescence. Approximately a half of them (58/(101+7) ≈ 0.54) belong to the actin cytoskeletal components that include NMMIIa (MYH9) and α-actinin-1 (ACTN1). These results seem to be consistent with the observations that SFs become large with RS, but we expected this first comprehensive analysis on the composition of SFs might include undiscovered proteins that alter the size of individual SFs. We then focused on elongation factor eEF2 as it exhibited the most significant upregulation with RS among the detected proteome that covers the diverse biological functions (eEF2, GSTP1, CD44, ITGB1, and THBS1; Fig. 4).

We found that eEF2 is condensed in SFs of cells undergoing RS (Fig. 5). Moreover, eEF2 was found to be critical to the RS-driven thickening (Fig. 6) and maturation of SFs (Fig. 7). There is a previous report, in which another elongation factor eEF1 was shown, surprisingly, to be involved in bundling the actin filaments to regulate the morphology of yeast cells independent of the mRNA translation function (Gross and Kinzy, 2005; Hamey and Wilkins, 2018). It remains unknown if eEF2 also possesses such similar actin bundling functions in fibroblasts in addition to its well-characterized translation activity; but, if this holds true, our results may reflect that SFs are stabilized in senescent human fibroblasts not only just by increasing the abundance of conventional actin bundling proteins of NMMIIa and α-actinin-1 but also by upregulating eEF2 as an additional actin cross-linker to further mature the macromolecular structure of SFs, while modulating the independent mRNA translation function. Determining the availability of eEF2 in actin bundling will be the subject of future investigation.

In recent studies, eEF2 has increasingly been implicated in neurosynaptic plasticity (Verpelli *et al*., 2010), NF-κB-associated innate immune response network (Bianco *et al*., 2019), and tumor progression as an oncogene (Oji *et al*., 2014; Sun *et al*., 2015; Rong *et al*., 2020). While the relevance of eEF2 to the cytoskeleton remains poorly elucidated, these discoveries may suggest its role in cytoskeletal regulation. In fact, close interactions between the actin cytoskeleton and protein translational machineries have been suggested (Kim and Coulombe, 2010; Silva *et al*., 2016; Simpson *et al*., 2020). Given this situation, our findings shed a new light on the molecular basis of the actin-based structure SFs and how their proteome is modulated to cope with senescence progression.

## Materials and methods

### Cell culture and transfection

Human foreskin fibroblasts HFF-1 (ATCC) were cultured with DMEM (High Glucose) including l-glutamine and phenol red (Wako) supplemented with 15% FBS (Sigma-Aldrich) and 1% penicillin-streptomycin solution (Wako) in a 5% CO_2_ incubator at 37°C. Cells were transfected with plasmids or siRNAs using Lipofectamine LTX Reagent with PLUS Reagent (Thermo Fisher Scientific) or RNAiMAX (Thermo Fisher Scientific), respectively, according to the manufacturer’s instructions. For passaging to induce RS, cells were seeded in polystyrene tissue culture flasks at an ~20% density and treated with trypsin at 80–90% confluency; consequently, it took about 80 days in total to obtain P33 cells.

### Plasmids and siRNAs

Plasmids encoding human MLC gene combined with EGFP were constructed in our group previously (Huang *et al*., 2021). The cDNAs encoding human elongation factor eEF2 were amplified by PCR using Q5 High-Fidelity 2X Master Mix (NEB) with primers (forward primer: GCGAAGCTTCGATGGTGAACTTCACGGT; reverse primer: GCGGCGAATTCCTACAATTTGTCCAGGAAG) and inserted into the pEGFP vector. The siRNAs targeting eEF2 and negative control siRNAs (s4991 and s4492, Thermo Fisher Scientific) were transfected to cells as described above.

### Western blotting

Cells were lysed in RIPA buffer (50 mM Tris-HCl, 100 mM NaCl, 1% NP-40, 1% sodium deoxycholate, 1 mM Na3VO4, 1 mM NaF, 1 mM dithiothreitol, 1 mM phenylmethylsulfonyl fluoride, 1μg/ml pepstatin, and 1 μg/ml leupeptin; pH 7.4) and centrifuged at 15000×g for 30 min to collect the supernatant. Protein concentration was measured by Pierce BCA Protein Assay Kit (Thermo Fisher Scientific). Proteins were fractionated in 10% gradient acrylamide gels (Bio-rad). Following SDS-PAGE, samples in gels were transferred onto polyvinylidene fluoride membranes (0.45 μm, Wako). After blocking in 5% BSA for 1 hour at room temperature, primary antibodies were used and detected using HRP-conjugated anti-rabbit or anti-mouse secondary antibodies (Bio-rad). The target bands were stained with Immobilon Western Kit (Millipore, Burlington, USA) and analyzed with ImageLab software (Bio-rad).

### Wound healing and proliferation assays

Cell migration was evaluated by performing wound healing assay. The healing rate was quantified 12 hours after a linear wounding was made on confluent cells by using a pipette tip. Cell proliferation was evaluated by initially incubating cells with EdU regent for 36 hours and then by staining them using Click-iT EdU Cell Proliferation Kit (Thermo Fisher Scientific).

### SA-β-gal staining

Cells cultured on 6-well plates were stained using SA-β-gal activity assay kit (Cell Signaling Technology) according to the manufacturer’s instruction. The staining images were taken using a microscope (IX73, Olympus) and analyzed with ImageJ software (NIH).

### Immunofluorescence

Cells cultured on glass-bottom dishes were fixed with 4% paraformaldehyde in phosphate-buffered saline (Wako) at 37°C for 30 min, permeabilized with 0.1% Triton X-100 for 15 min, blocked with 5% normal goat serum for 1 hour, and incubated with the primary antibody for 1.5 hours and then hybridized with appropriate secondary antibodies (Thermo Fischer Scientific) for 1 hour at room temperature. The cell nucleus and actin filaments were stained with Hoechst 33342 (Thermo Fischer Scientific) and fluorescently-labeled phalloidin (Thermo Fischer Scientific), respectively. Images were acquired using a microscope (IX73, Olympus) or a confocal laser scanning microscope (FV1000, Olympus) equipped with a UPlan Apo 60x oil objective lens (NA = 1.42) and analyzed using ImageJ software.

### Isolation of SFs from cells

Cells were cultured to reach approximately 80% confluency on 60-mm-diameter polystyrene culture dishes. Cells were hypotonically shocked with a low-ionic strength solution (2.5mM triethanolamine, 1 mM dithiothreitol, 1μg/ml pepstatin, and 1 μg/ml leupeptin in ultra-pure water) for 2 min and then were subjected to fluid shearing forces by repeated pipetting in the low-ionic solution for 30 sec. The buffer was collected by Centrifugal Filters Kit (Amicon Ultra) to serve as the cell body lysates for western blot. Ventral SFs remaining bound to the dish were incubated with 0.05% Triton X-100 in a cytoskeleton stabilizing buffer (20 mM Imidazole, 2.2 mM MgCl_2_, 2 mM EGTA, 13.3 mM KCl, 1 mM dithiothreitol, 1 μg/ml pepstatin, and 1 μg/ml leupeptin; pH 7.3 (Matsui *et al*., 2011; Deguchi *et al*., 2012; Okamoto *et al*., 2020)) for 1 min and then were rinsed 3 times with the same buffer free of Triton X-100. The extracted SFs were collected in RIPA buffer on ice.

### Mass spectrometry-based shotgun proteomics

Cells at P2 and P33 were lysed with non-denature lysis buffer (50 mM Tris, 2% sodium deoxycholate, 1 mM DTT, 10 μg/ml leupeptin,10 μg/ml pepstatin, and 1 mM PMSF; pH 8.0). The sample lysates were sequenced by mass spectrometry at the Center for Medical Innovation and Translational Research (COMIT) of Osaka University Medical School using UltiMate 3000 RSLCnano system coupled with Q-Exactive mass spectrometer (Thermo Fisher Scientific) to distinguish between P2 and P33 by identifying protein expression differences. The false discovery rate was set to 1.4% at peptide level. Functional analysis was performed using Scaffold 4 software (Proteome Software).

### Fluorescence recovery after photobleaching

Cells expressing EGFP-MLC were cultured on a glass-bottom dish in a stage incubator (Tokai Hit). FRAP experiments were performed using a confocal microscope (FV1000, Olympus) with a 60x oil immersion objective lens on cells treated in advance with siRNAs for control or for eEF2 for 36 hours before the experiments. Photobleaching was induced using the 405/440-nm-wavelength lasers on individual MLC-labeled SFs, and images were taken for 4 min at a 5-sec-interval. The recovery curve was fitted to a single exponential function by the least-squares method, by which the steady-state value of the fluorescence intensity, i.e., mobile fraction, was computed to measure the instability of SFs.

### Statistical analysis

Statistical analysis was performed using Prism 8 (GraphPad Software), in which *p-* values were calculated using a one-way analysis of variance followed by Tukey’s test or two-tailed *t*-test for multiple comparisons. Data were shown as mean + standard deviation from more than three independent experiments. Statistical significance was set, compared to respective controls, as follows: *, *p* < 0.05, **, *p* < 0.01; and ***, *p* < 0.001.

## Acknowledgments

SL and NK are supported by Chinese Scholarship Council (CSC) Scholarship. This work was supported in part by JSPS KAKENHI Grants (18H03518 and 19K22967).

## Dada availability statement

The data that support the findings of this study are available from the corresponding author upon reasonable request.

## Supplementary materials

**Figure S1:**
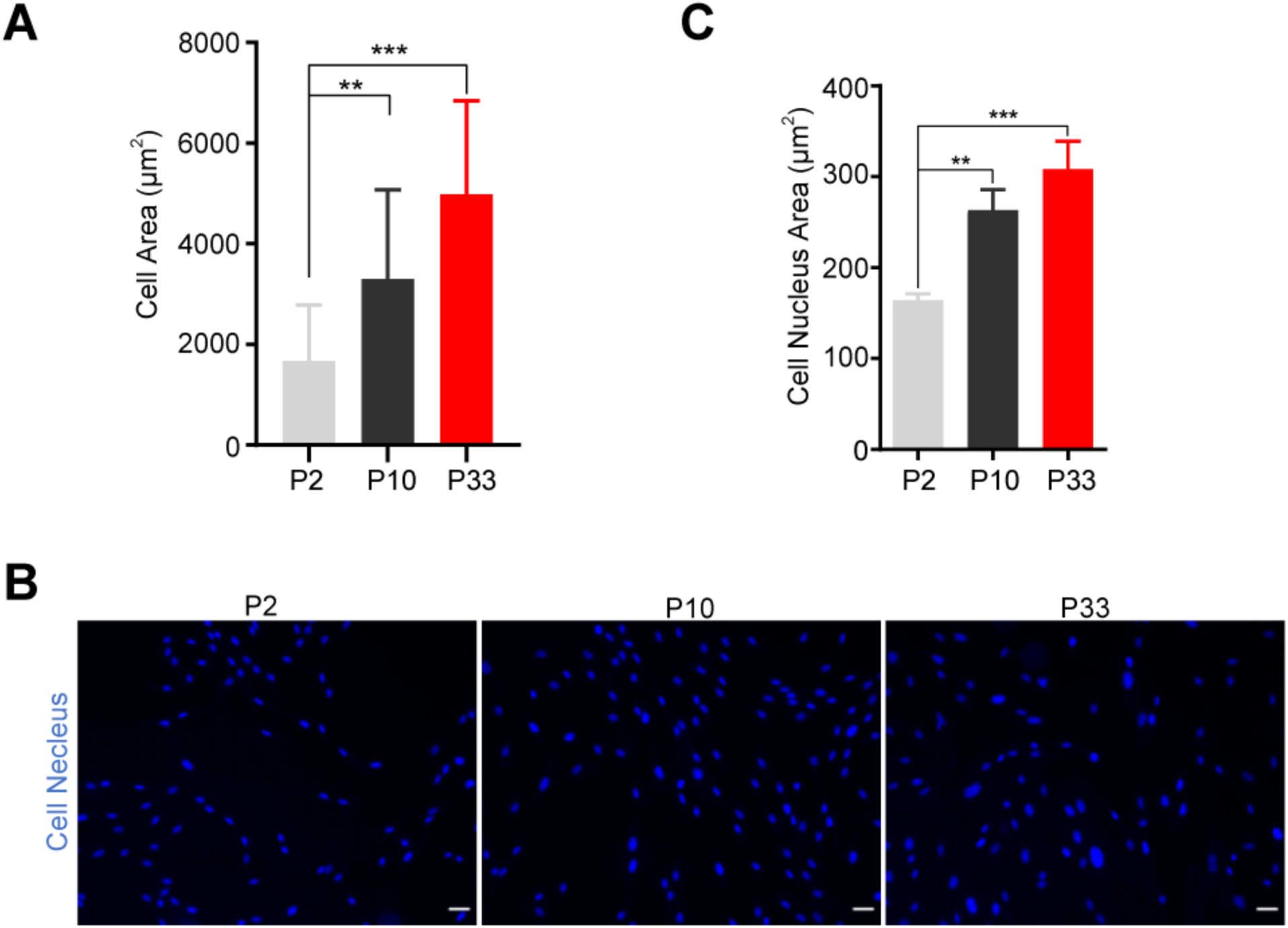
Morphological responses to RS. (A) Quantification of the cell images (as shown in Fig. 2A) shows that cell area is increased with RS. (B) The nuclei are stained with Hoechst 33342 in cells at different senescence levels. (C) Quantification shows that the nuclear area is increased with RS. Scale, 100 μm.

**Figure S2:**
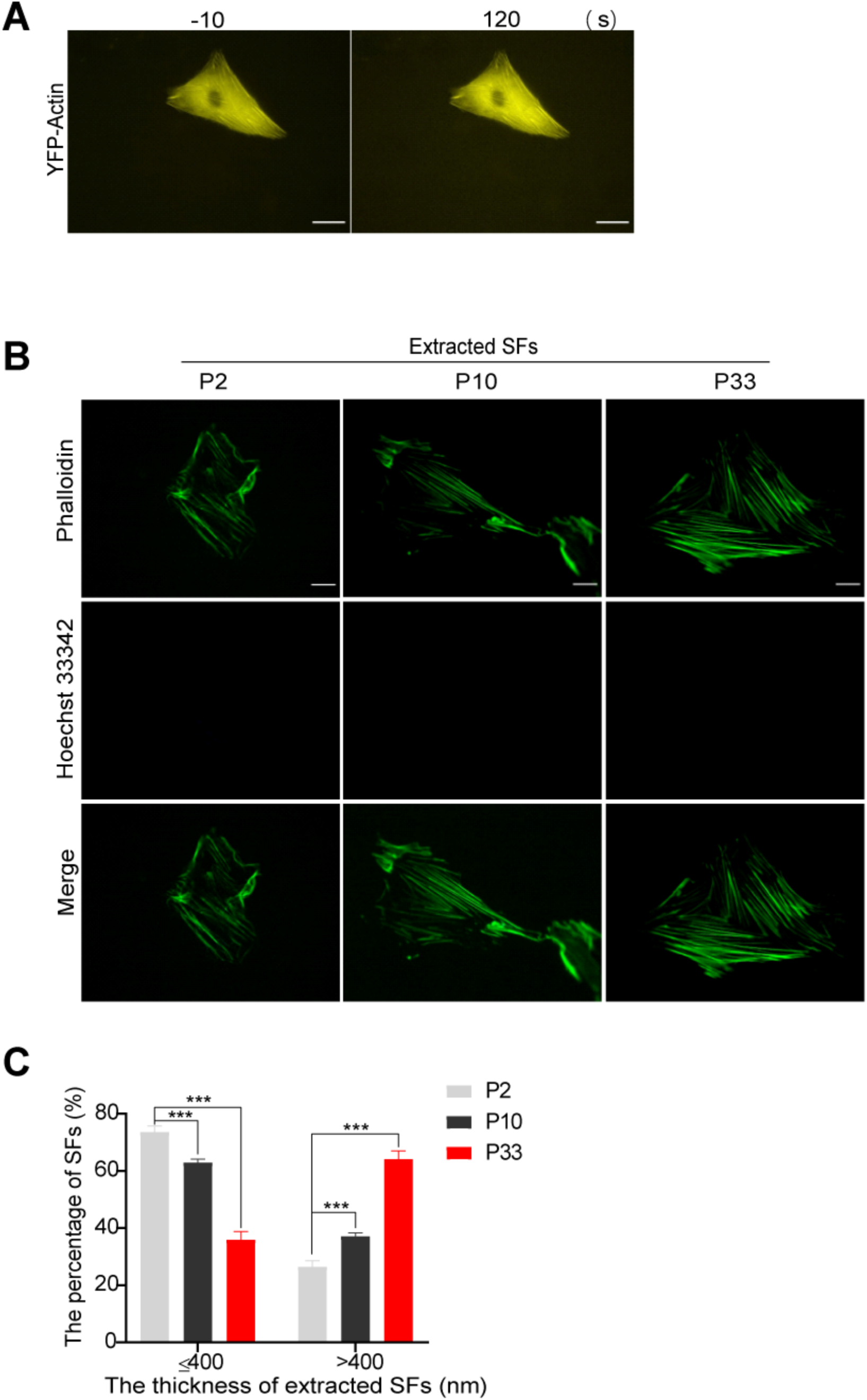
SFs remain unchanged in morphology with the extraction treatment. (A) Images of YFP-actin in living HFF-1 cells exposed to the low-ionic strength solution show that the morphology of SFs is not affected by the treatment. (B) The morphology of extracted SFs (actin filaments stained with phalloidin) remaining bound to the substrate after the treatment with Triton X-100 but before the collection. Note that the nuclei are removed in the extracted samples. (C) Quantification shows that the extracted SFs maintain increased thickness similar to that of intact SFs. Scale, 20 μm.

Figure S3: Classification of SF-associated proteins identified in the proteomic analysis according to the GeneCards database. The figure in parenthesis shows the number of proteins classified in each group: in total, 263 proteins are detected in the isolated SFs; 73 are described in the database to be associated with the cytoskeleton, and among them, 15 are typical for intermediate filaments or microtubules, and the remaining 58 are typical for actin filaments. Further categorizations are done according to the database and shown in different colors. The groups are also categorized according to the measured FC, i.e., a ratio of the expression at P33 to that at P2.

## Reference

Bianco, C., Thompson, L., and Mohr, I. (2019). Repression of eEF2K transcription by NF-kappaB tunes translation elongation to inflammation and dsDNA-sensing. Proc Natl Acad Sci U S A 116, 22583–22590.

Braga, V.M., Machesky, L.M., Hall, A., and Hotchin, N.A. (1997). The small GTPases Rho and Rac are required for the establishment of cadherin-dependent cell-cell contacts. J Cell Biol 137, 1421–1431.

Burridge, K., and Wittchen, E.S. (2013). The tension mounts: stress fibers as force-generating mechanotransducers. J Cell Biol 200, 9–19.

Chen, C.S., Tan, J., and Tien, J. (2004). Mechanotransduction at cell-matrix and cell-cell contacts. Annu Rev Biomed Eng 6, 275–302.

Chen, Q.M., Tu, V.C., Catania, J., Burton, M., Toussaint, O., and Dilley, T. (2000). Involvement of Rb family proteins, focal adhesion proteins and protein synthesis in senescent morphogenesis induced by hydrogen peroxide. J Cell Sci 113 (Pt 22), 4087–4097.

Chien, S. (2007). Mechanotransduction and endothelial cell homeostasis: the wisdom of the cell. Am J Physiol Heart Circ Physiol 292, H1209–1224.

Choi, J., Grosely, R., Prabhakar, A., Lapointe, C.P., Wang, J.F., and Puglisi, J.D. (2018). How Messenger RNA and Nascent Chain Sequences Regulate Translation Elongation. Annu Rev Biochem 87, 421–449.

Cristofalo, V.J., Lorenzini, A., Allen, R.G., Torres, C., and Tresini, M. (2004). Replicative senescence: a critical review. Mech Ageing Dev 125, 827–848.

Deguchi, S., Matsui, T.S., Komatsu, D., and Sato, M. (2012). Contraction of Stress Fibers Extracted from Smooth Muscle Cells: Effects of Varying Ionic Strength. Journal of Biomechanical Science and Engineering 7, 388–398.

Deguchi, S., Ohashi, T., and Sato, M. (2006). Tensile properties of single stress fibers isolated from cultured vascular smooth muscle cells. J Biomech 39, 2603–2610.

Goldstein, S. (1990). Replicative senescence: the human fibroblast comes of age. Science 249, 1129–1133.

Gross, S.R., and Kinzy, T.G. (2005). Translation elongation factor 1A is essential for regulation of the actin cytoskeleton and cell morphology. Nat Struct Mol Biol 12, 772–778.

Hamey, J.J., and Wilkins, M.R. (2018). Methylation of Elongation Factor 1A: Where, Who, and Why? Trends Biochem Sci 43, 211–223.

Hayflick, L. (1965). The Limited in Vitro Lifetime of Human Diploid Cell Strains. Exp Cell Res 37, 614–636.

Hirata, H., Gupta, M., Vedula, S.R., Lim, C.T., Ladoux, B., and Sokabe, M. (2015). Actomyosin bundles serve as a tension sensor and a platform for ERK activation. EMBO Rep 16, 250–257.

Hotulainen, P., and Lappalainen, P. (2006). Stress fibers are generated by two distinct actin assembly mechanisms in motile cells. J Cell Biol 173, 383–394.

Huang, W., Matsui, T.S., Saito, T., Kuragano, M., Takahashi, M., Kawahara, T., Sato, M., and Deguchi, S. (2021). Mechanosensitive myosin II but not cofilin primarily contributes to cyclic cell stretch-induced selective disassembly of actin stress fibers. Am J Physiol Cell Physiol.

Hwang, E.S., Yoon, G., and Kang, H.T. (2009). A comparative analysis of the cell biology of senescence and aging. Cell Mol Life Sci 66, 2503–2524.

Jalal, S., Shi, S., Acharya, V., Huang, R.Y., Viasnoff, V., Bershadsky, A.D., and Tee, Y.H. (2019). Actin cytoskeleton self-organization in single epithelial cells and fibroblasts under isotropic confinement. J Cell Sci 132.

Kang, N., Matsui, T.S., and Deguchi, S. (2021). Statistical profiling reveals correlations between the cell response to and the primary structure of Rho-GAPs. Cytoskeleton (Hoboken).

Kang, N., Matsui, T.S., Liu, S., Fujiwara, S., and Deguchi, S. (2020). Comprehensive analysis on the whole Rho-GAP family reveals that ARHGAP4 suppresses EMT in epithelial cells under negative regulation by Septin9. FASEB J 34, 8326–8340.

Kaunas, R., and Deguchi, S. (2011). Multiple Roles for Myosin II in Tensional Homeostasis Under Mechanical Loading. Cell Mol Bioeng 4, 182–191.

Kaunas, R., Usami, S., and Chien, S. (2006). Regulation of stretch-induced JNK activation by stress fiber orientation. Cell Signal 18, 1924–1931.

Kim, S., and Coulombe, P.A. (2010). Emerging role for the cytoskeleton as an organizer and regulator of translation. Nat Rev Mol Cell Biol 11, 75–81.

Kreis, T.E., and Birchmeier, W. (1980). Stress fiber sarcomeres of fibroblasts are contractile. Cell 22, 555–561.

Kumar, S., Maxwell, I.Z., Heisterkamp, A., Polte, T.R., Lele, T.P., Salanga, M., Mazur, E., and Ingber, D.E. (2006). Viscoelastic retraction of single living stress fibers and its impact on cell shape, cytoskeletal organization, and extracellular matrix mechanics. Biophys J 90, 3762–3773.

Kuo, J.C., Han, X., Hsiao, C.T., Yates, J.R., 3rd, and Waterman, C.M. (2011). Analysis of the myosin-II-responsive focal adhesion proteome reveals a role for beta-Pix in negative regulation of focal adhesion maturation. Nat Cell Biol 13, 383–393.

Levine, A.J., and Oren, M. (2009). The first 30 years of p53: growing ever more complex. Nat Rev Cancer 9, 749–758.

Lin, Y.H., Zhen, Y.Y., Chien, K.Y., Lee, I.C., Lin, W.C., Chen, M.Y., and Pai, L.M. (2017). LIMCH1 regulates nonmuscle myosin-II activity and suppresses cell migration. Mol Biol Cell 28, 1054–1065.

Livne, A., and Geiger, B. (2016). The inner workings of stress fibers - from contractile machinery to focal adhesions and back. J Cell Sci 129, 1293–1304.

Matsui, T.S., Kaunas, R., Kanzaki, M., Sato, M., and Deguchi, S. (2011). Non-muscle myosin II induces disassembly of actin stress fibres independently of myosin light chain dephosphorylation. Interface Focus 1, 754–766.

McHugh, D., and Gil, J. (2018). Senescence and aging: Causes, consequences, and therapeutic avenues. J Cell Biol 217, 65–77.

Meng, W., and Takeichi, M. (2009). Adherens junction: molecular architecture and regulation. Cold Spring Harb Perspect Biol 1, a002899.

Muller, M. (2009). Cellular senescence: molecular mechanisms, in vivo significance, and redox considerations. Antioxid Redox Signal 11, 59–98.

Nishio, K., and Inoue, A. (2005). Senescence-associated alterations of cytoskeleton: extraordinary production of vimentin that anchors cytoplasmic p53 in senescent human fibroblasts. Histochem Cell Biol 123, 263–273.

Oji, Y., Tatsumi, N., Fukuda, M., Nakatsuka, S., Aoyagi, S., Hirata, E., Nanchi, I., Fujiki, F., Nakajima, H., Yamamoto, Y., Shibata, S., Nakamura, M., Hasegawa, K., Takagi, S., Fukuda, I., Hoshikawa, T., Murakami, Y., Mori, M., Inoue, M., Naka, T., Tomonaga, T., Shimizu, Y., Nakagawa, M., Hasegawa, J., Nezu, R., Inohara, H., Izumoto, S., Nonomura, N., Yoshimine, T., Okumura, M., Morii, E., Maeda, H., Nishida, S., Hosen, N., Tsuboi, A., Oka, Y., and Sugiyama, H. (2014). The translation elongation factor eEF2 is a novel tumorassociated antigen overexpressed in various types of cancers. Int J Oncol 44, 1461–1469.

Okamoto, T., Matsui, T.S., Ohishi, T., and Deguchi, S. (2020). Helical structure of actin stress fibers and its possible contribution to inducing their direction-selective disassembly upon cell shortening. Biomech Model Mechanobiol 19, 543–555.

Rong, F., Liu, L., Zou, C., Zeng, J., and Xu, Y.S. (2020). MALAT1 Promotes Cell Tumorigenicity Through Regulating miR-515-5p/EEF2 Axis in Non-Small Cell Lung Cancer. Cancer Manag Res 12, 7691–7701.

Silva, R.C., Sattlegger, E., and Castilho, B.A. (2016). Perturbations in actin dynamics reconfigure protein complexes that modulate GCN2 activity and promote an eIF2 response. J Cell Sci 129, 4521–4533.

Simpson, L.J., Reader, J.S., and Tzima, E. (2020). Mechanical Forces and Their Effect on the Ribosome and Protein Translation Machinery. Cells 9.

Sun, W., Wei, X., Niu, A., Ma, X., Li, J.J., and Gao, D. (2015). Enhanced anti-colon cancer immune responses with modified eEF2-derived peptides. Cancer Lett 369, 112–123.

Tojkander, S., Gateva, G., and Lappalainen, P. (2012). Actin stress fibers--assembly, dynamics and biological roles. J Cell Sci 125, 1855–1864.

van Deursen, J.M. (2014). The role of senescent cells in ageing. Nature 509, 439–446.

Verpelli, C., Piccoli, G., Zanchi, A., Gardoni, F., Huang, K., Brambilla, D., Di Luca, M., Battaglioli, E., and Sala, C. (2010). Synaptic Activity Controls Dendritic Spine Morphology by Modulating eEF2-Dependent BDNF Synthesis. Journal of Neuroscience 30, 5830–5842.

Wei, F., Xu, X., Zhang, C., Liao, Y., Ji, B., and Wang, N. (2020). Stress fiber anisotropy contributes to force-mode dependent chromatin stretching and gene upregulation in living cells. Nat Commun 11, 4902.

